# Phagocytosis repurposed: infection strategies in a global marine diatom-parasite interaction

**DOI:** 10.1101/2025.07.29.667418

**Authors:** Varsha Mathur, Nicholas A.T. Irwin, Luis Javier Galindo, Elisabet Alacid, Michael Cunliffe, Sunju Kim, Ulrich Technau, Thomas A Richards

**Affiliations:** Department of Biology, University of Oxford, Oxford, United Kingdom; Faculty of Life Sciences, University of Vienna, Vienna, Austria; Gregor Mendel Institute (GMI), Austrian Academy of Sciences, Vienna, Austria; Institute of Water Research, University of Granada, Granada, Spain; Department of Ecology, University of Granada, Granada, Spain; Centre d’Estudis Avançats de Blanes (CEAB-CSIC), Blanes, Spain; School of Biological and Marine Sciences, University of Plymouth, United Kingdom; Department of Oceanography, Pukyong National University, Busan 48513, Korea

## Abstract

Microbial host–parasite interactions shape global biogeochemical cycles, yet their cellular and evolutionary mechanisms remain poorly understood. Here, we developed a novel model pathosystem to explore the interaction between the globally distributed eukaryotic microparasite, *Pirsonia,* and its host, the bloom-forming diatom, *Coscinodiscus*. Culture conditions recapitulated *Pirsonia*’s rapid life cycle *in vitro* and live-cell imaging enabled quantitative assessments of infection dynamics and host mortality. To investigate the genetic basis of this interaction, we assembled the *Pirsonia* genome and identified unique genetic innovations relative to their oomycete relatives. Time-resolved dual RNA-Seq enabled tracking of parasite gene expression through the infection cycle and revealed an upregulation of an expanded multi-gene family of integrin-like proteins in *Pirsonia* zoospores, analogous to the variable surface proteins in other parasitic lineages. Genes upregulated during infection were associated with cytoskeletal dynamics and drug inhibition assays confirmed that actin is required for host infection, consistent with the presence of parasitic pseudopodia on infected hosts. The dependence on both actin-based phagocytic mechanisms and parasitic-like surface proteins highlights *Pirsonia*’s intermediate state between predator and parasite, providing new insights into the evolution of parasite strategies and the complex cellular interactions controlling parasitic shunts in marine trophic networks.

## MAIN

The ocean contains a diverse array of microorganisms including bacteria, archaea, viruses, fungi, and microbial eukaryotes^1–3^. These species form the base of marine trophic webs and contribute to the biogeochemical cycling of vital nutrients, particularly fixed carbon and nitrogen^4^. Eukaryotic phytoplankton, such as diatoms and coccolithophores, account for 1–2% of the Earth’s primary producer biomass yet are responsible for 40% of global annual carbon-based photosynthesis^5^. This is in part driven by the formation of algal blooms, which occur following rapid proliferation^6^. The ecological implications of these blooms are manifold, as they can result in harmful toxin production^7^ but also form hotspots for microbial interactions that have large-scale impacts on marine metabolic flux and ultimately global chemical cycles^8,9^. Accordingly, understanding the ecological and cellular interactions governing bloom dynamics are important for understanding ocean ecosystems.

Although nutrient availability and climatic conditions can drive bloom formation, pathogen infections play a major role in bloom collapse^10^. In particular, viral infections have been shown to impact marine algal bloom dynamics and carbon flow through a process termed the “viral shunt”^11,12^. Similarly, eukaryotic microparasites are significant mortality agents, regulating diverse phytoplankton populations such as dinoflagellates and diatoms^13–18^, and are consistently found in high abundances in metabarcoding surveys of ocean environments^19^. Despite this, the impact of microparasites on planktonic networks and biogeochemistry are rarely considered^2^. Establishing experimentally tractable pathosystems for understanding these relationships has been challenging due to the complexities of parasite isolation and host-parasite co–culture maintenance. Nonetheless, a few model pathosystems have been established shedding light on the metabolic and transcriptomic reprogramming during parasitic infections^20–22^. In addition, single-cell transcriptomics and metabarcoding approaches have also facilitated studies into uncultured marine parasites^23^. However, developing diverse and tractable host-parasite model systems remains critical for gaining genomic and cellular insights into the diversity of these interactions.

To address this, we sought to establish a new understanding of a marine model pathosystem tractable for laboratory study; a system involving the globally distributed parasite, *Pirsonia diadema* and its diatom hosts, *Coscinodiscus. Pirsonia* is a relative of the Pseudofungi, a group including the oomycetes, and exhibits a parasitoid life strategy, where it exploits the host for its development, ultimately leading to host mortality^24–26^. *P. diadema* is a host-specific parasite of the centric diatom genus *Coscinodiscus* which forms seasonal blooms and is protected by a silica cell wall, known as the frustule^27^. To better understand the molecular and cellular dynamics underpinning this interaction, we established long-term co-cultures of *Pirsonia* and *Coscinodiscus*. Live-cell imaging enabled quantitative assessments of infection progression and genomic and transcriptomic analyses revealed lineage-specific innovations in parasitic surface proteins and cytoskeletal regulators critical for infection. These data highlight *Pisonia’s* intermediate state between parasite and predator, revealing new insights into parasitic evolution and the mechanisms controlling a key microbial interaction in the ocean.

### Establishing a new ecologically relevant pathosystem

To investigate the *Pirsonia-Coscinodiscus* interactions, we first aimed to establish a stable co-culture of these species. To do this, we performed single-cell isolations to establish clonal cultures from natural plankton samples collected from the Western English Channel, UK, and the Yongho Bay of Busan, Republic of Korea. We observed the life cycle and cellular morphology of *P. diadema* using light microscopy and scanning electron microscopy (SEM) (Fig. 1a-c). In culture, we observed the motile, bi-flagellate zoospores of *Pirsonia* attached to the diatom frustule, specifically on the sites of diatom pores, known as rimoportulae (Fig. 1c, white arrow) where they penetrated the host intracellular environment. The zoospores would subsequently lose their flagella and differentiate into auxosomes, many of which were frequently seen on a single host cell (Fig. 1b). During infection, the diatom protoplast was remodelled into digestive compartments known as trophosomes, while the auxosome remained on the diatom surface, growing and dividing. Following host death, the auxosomes detached and formed flagellate mother cells (FMCs), which exhibit motility and possessed a single flagellum, leaving behind an empty diatom frustule. The FMCs would eventually divide and differentiate into numerous bi-flagellated, fast-swimming, infective zoospores, completing the previously described parasitoid life cycle^25^.

**Figure 1:**
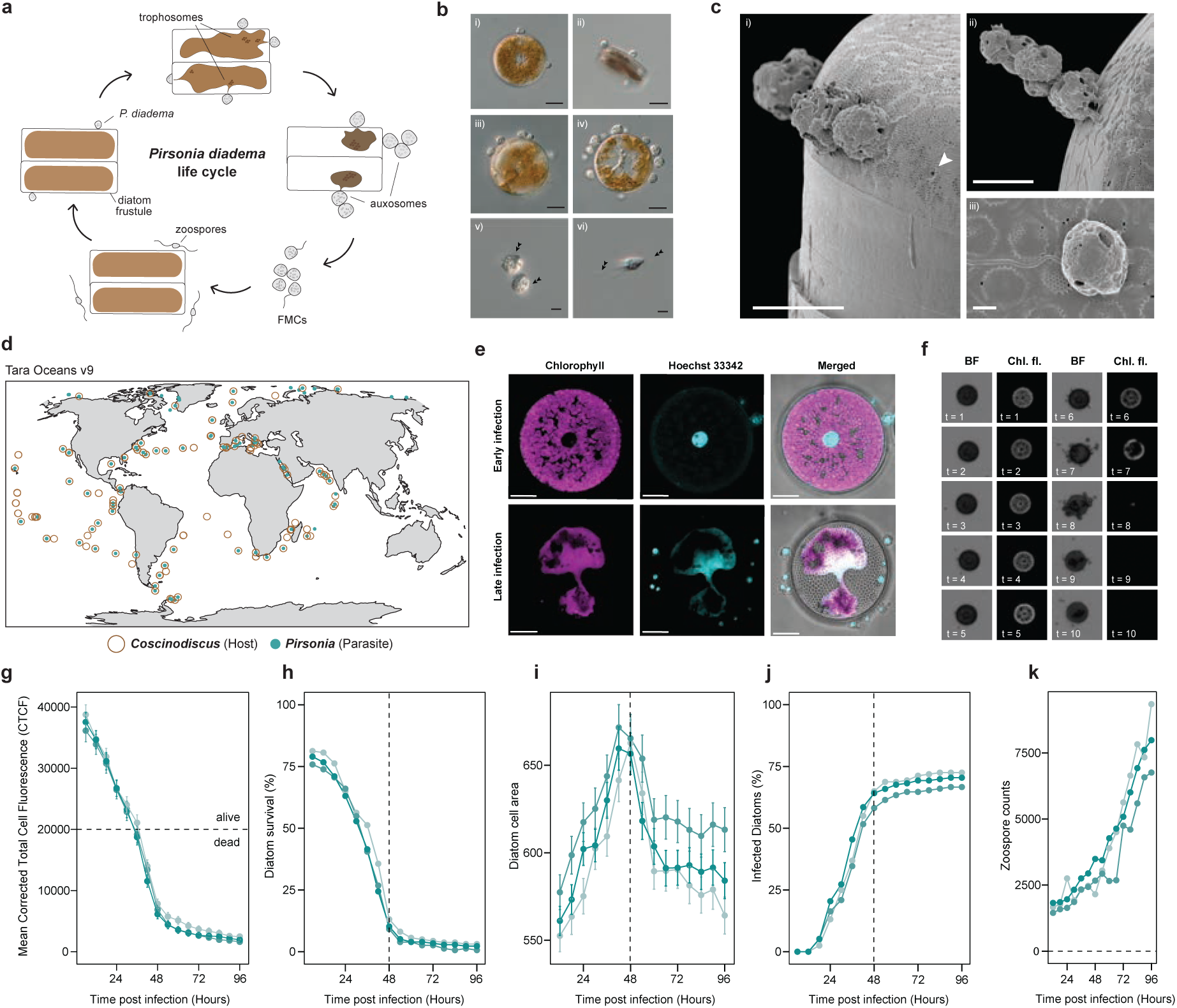
Microscopic characterization of the *Pirsonia*-*Coscinodiscus* model system. **a,** Schematic of the developmental stages and life cycle of *P. diadema* with *Coscinodiscus* shown in girdle view. **b,** Light micrographs of *P. diadema* infecting *C. radiatus* (top two panels) and two distinct flagellate stages: flagellate mother cells (FMCs) and zoospores (bottom panel). Scale bars, 20 μm (i–iv), 5 μm (v-vi). **c,** Frustule surface host-parasite interaction visualised by scanning electron microscopy. White arrow indicates rimoportulae (sites of parasitoid attachment). Scale bars, 5 μm (i–ii), 1 μm (iii). **d,** Geographical distribution of rDNA 18S V9 environmental OTUs (Operational Taxonomic Units) belonging to the genus *Pirsonia* and genus *Coscinodiscus* in the Tara Oceans Expedition. **e,** Fluorescence microscopy of early (top panel) and late (bottom panel) infections. Chlorophyll autofluorescence (Chl. Fl) -purple and Hoechst 33342 DNA - light blue staining of *C. radiatus* infected with *P. diadema*. Scale bars, 10 μm. **f,** Automated live-cell high-content screening (HCS) of *P. diadema* infection. Brightfield (BF) and Chl. Fl micrographs of an individual cell tracked every 6 hours over a 4-day period. t = timepoint. **g-k,** Line graphs quantifying infection dynamics using HCS to track the fate of individual diatom cells (n = 489) when infected with *P. diadema*. Error bars indicate standard deviation, and colors represent biological replicates. **g,** Mean corrected total cell fluorescence (CTCF) of infected diatoms over time. Horizontal dashed line indicates the Chl. Fl. threshold for cell viability. **h,** Diatom survival curve calculated based on CTCF levels. Vertical dashed line marks the time point where widespread cell death occurred. **i,** Diatom cell area over time post-infection. The vertical dashed line marks the time point where cell area peaks. **j,** Percentage of infected diatoms over time based on area, reaching a plateau after ∼48 hours (vertical dashed line). **k,** Line graph displaying zoospores numbers over time post-infection.

Given the establishment and observation of the complete pathogenic life cycle in the laboratory, we next sought to assess the ecological relevance of the interaction using environmental sequencing data. We analysed the 18S rDNA V9 metabarcoding dataset from the Tara Oceans Expedition^28^ and identified OTUs (operational taxonomic units) corresponding to both *Pirsonia* and *Coscinodiscus* across global oceans (Fig. 1d, Fig. S1, Table S1). Both organisms have widespread geographic ranges, with *Pirsonia* consistently co-occurring with *Coscinodiscus* except at a few polar sites, consistent with its obligate dependence on its diatom host. Their broad and overlapping distributions across diverse marine environments underscores the ecological significance of this interaction in global planktonic systems.

### The temporal dynamics of the *Pirsonia-Coscinodiscus* infection cycle

To gain deeper insights into the cellular changes occurring during diatom infection, we used fluorescence confocal microscopy (Fig 1e). During the early stages of infection, the subcellular integrity of the diatom protoplast remains intact, with the nucleus positioned in the centre of the cell surrounded by a dense network of chloroplasts exhibiting strong chlorophyll autofluorescence (Fig. 1e, top row). However, in the later stages of infection, we observed a marked decrease in chlorophyll autofluorescence and DNA staining, as the diatom protoplasm is degraded by *Pirsonia* trophosomes (Fig. 1e, bottom row).

Leveraging the marked loss of chlorophyll autofluorescence as a proxy for host viability and infection progression, we performed a quantitative analysis of infection dynamics using an automated live-cell high-content screener (HCS). This approach allowed for the tracking and measurement of individual diatom cells (*n* = 489) following exposure to *Pirsonia* zoospores, with imaging conducted every six hours over a four-day period (Fig. 1f, Supplemental video 1).

A decline in CTCF was evident within 12 hours post-infection and diatoms were classified as dead once their CTCF dropped below 50% of their initial value (Fig. 1f-h). Using this threshold, diatom survival curves showed that most host cells had died within 48 hours of infection (Fig. 1h). To further quantify infection progression, we also measured changes in host cell area over time which could be attributed to the accumulation of auxospores on the host surface, given normal cell size limitations imposed by the frustule (Fig. 1b,f,i). Indeed, diatom death at 48 hours coincided with a peak in cell area (Fig. 1i). This phenotyping assay enabled precise inference of infection establishment timing (Fig. 1j), calculated as the point at which cell size increased by 10% of the initial value. *Pirsonia* infections occurred rapidly, and synchronously, with all diatoms infected between 24- and 48-hours post-infection. By 96 hours, all diatoms had died, while zoospore numbers increased exponentially following differentiation from the released auxospores (Fig. 1k). Overall, our live-cell timelapse imaging enabled quantitative, single-cell tracking of *Pirsonia* infections, revealing a life cycle defined by parasite establishment and development, host consumption and mortality, and zoospore proliferation, all within two days. These observed infection dynamics highlight the capacity of *Pirsonia* to rapidly impact diatom population structure.

### The *P. diadema* genome reveals lineage specific innovations distinct from other pseudofungal parasites

To better understand the genetic basis for *Pirsonia’s* rapid and highly virulent life cycle, we sought to characterize its genomic repertoire by generating a *de novo* draft genome using PacBio HiFi long-read sequencing. After performing stringent contamination removal (see Material and Methods), we assembled a genome of 5,155 scaffolds spanning 171 Mb, with a scaffold N50 of 42.1 Kb using Hifiasm^29^ (Fig 2a). Gene annotation identified 35,947 protein-coding genes with a BUSCO completeness score of 78% (Fig 2d). Together, these genomic statistics confirm the quality of the *P. diadema* genome, which is to our knowledge, the first genome sequenced to date from the Bigryomonada subphylum.

**Figure 2:**
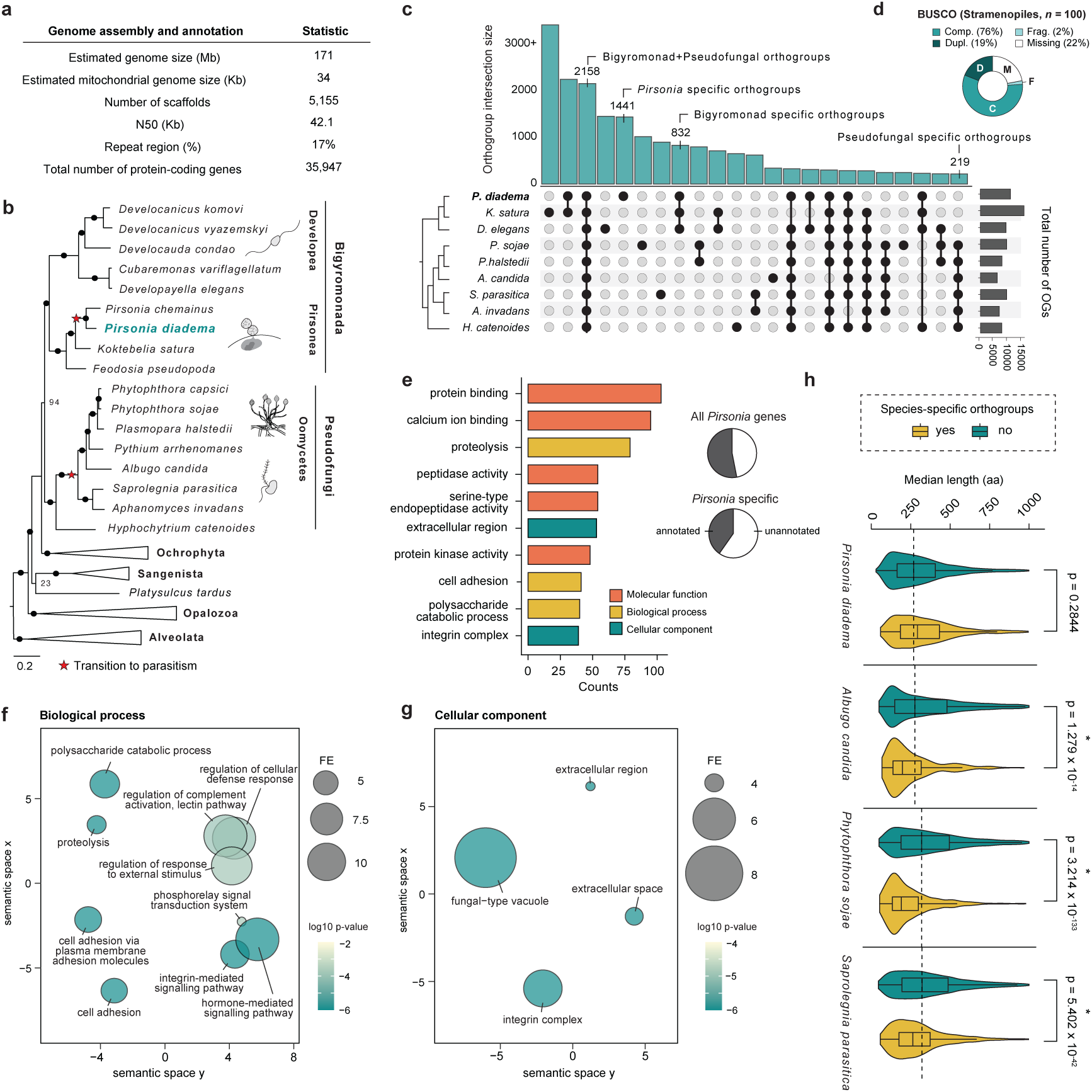
Comparative genomics of *P. diadema*. **a**, Summary of the *P. diadema* genome assembly and annotation statistics. **b,** Maximum likelihood tree of stramenopiles (with alveolate outgroup) generated from 260 genes using LG+C40+F+G4 substitution model implemented in IQ-Tree. Support was obtained 100 standard non-parametric bootstraps (NPB) and 1,000 ultrafast bootstraps (UFB). Black circles represent NPB/UFB values greater than 95% and transitions to parasitism are indicated. **c,** Upset plot displaying the shared and unique orthogroups between across the Pseudofungi. Each row represents a pseudofungal species. Black circles and vertical lines between the rows represent the intersection of orthogroups. The cladogram (left) illustrates the phylogenetic relationships between species and the barplot (right) indicates the total number of orthogroups in each species. OGs = orthogroups. **d,** Donut chart displaying a summary of complete, fragmented, duplicated and missing BUSCO genes in *P. diadema* genome based on the stramenopile_odb10 lineage dataset. **e,** Barplot displaying the most frequent gene ontology (GO) terms found in the *P. diadema* genome annotations based on counts. The colors indicate whether the GO term is a molecular function (red), biological process (yellow), or cellular component (green). The pie charts (top) display the proportion of *P. diadema* orthogroups that have GO annotation and (bottom) the proportion of *P. diadema*-specific orthogroups that have GO annotations. **f, g,** Scatterplots displaying enriched GO biological process and cellular component terms in *P. diadema* orthogroups annotations. Semantic similarity was determined using REVIGO and statistical significance was assessed using two-sided permutation tests (*P* < 0.01, *n* = 10^7^). **h,** Violin plots comparing the median length of proteins in species-specific (yellow) and non-species-specific (teal) orthogroups in *P. diadema* and in three oomycete species. Horizontal dashed line marks the median length of non-species-specific orthogroups. P-values were assessed using two sample t-tests implemented in R and significance is denoted by an asterisk (*).

To confirm the phylogenetic placement of *Pirsonia diadema*, we generated a multi-gene phylogeny of Stramenopiles (Fig 2b, Table S2). As expected, our analyses placed *P. diadema* within the subphylum Bigyromonada, the sister group to the oomycetes, a group containing some of the most important agricultural parasites^30^, consistent with 18S rRNA gene phylogenies^24,26^. While oomycetes exhibit a fungal-like morphology and have been extensively sampled at the genomic level, bigyromonads comprise primarily of heterotrophic flagellates^31^. Interestingly, *Koktebelia,* a free-living eukaryovorous predator^31,32^, is the closest sister genus to *Pirsonia*, which is itself nested among other free-living bigyromonads. Overall, this phylogeny supports the idea that *Pirsonia* represents an independent transition to parasitism within the larger clade, in addition to at least three other distinct transitions within the oomycetes^33,34^. These findings underscore the multiple, convergent origins of parasitism across the Pseudofungi and Bigyromonada.

Given their parallel parasitic origins, we looked to compare the gene repertoires of *Pirsonia* with other parasitic and free-living members of the Pseudofungi. To do this, we clustered a representative set of pseudofungal proteomes into 98,614 orthologous groups using OrthoFinder^35^ (Fig. 2c, Table S3). Overall, bigyromonads exhibit a greater number of group-specific and species-specific orthogroups compared to the oomycetes, suggesting a higher level of divergence in their gene content and highlighting the limited sampling of bigyromonad diversity. *Pirsonia* shares the most orthogroups with the transcriptome of *K. satura*, as well as with other bigyromonad species. Additionally, *Pirsonia* possesses a substantial number of species-specific orthogroups (1,441). To explore the functions of these orthogroups, we examined their annotations and found that only 40% had functional assignments, indicative of a large proportion of predicted proteins with unknown functions (Fig. 2e). Nonetheless, the *Pirsonia*–specific annotated orthogroups were enriched in functions such as cell adhesion, cellular breakdown (proteolysis and polysaccharide catabolysis), signalling and regulation of cellular responses (Fig. 2e,f), processes which could be related to infectivity. In agreement, the cellular components such as extracellular space/region, including proteins involved in integrin complexes, were strongly enriched. This raises the hypothesis that these proteins may represent effector proteins secreted by *Pirsonia* or proteins found on the cell surface and involved in host interaction (Fig. 2g). In the oomycetes, effectors, such as proteins carrying an RxLR N-terminal motif^36^, are typically short and rapidly evolving. However, we found no significant reduction in the length of *Pirsonia*-specific proteins compared to the rest of the proteome (Fig. 1h). These differences may reflect the different evolutionary pressures imposed by host immune systems - specifically, between a single-celled diatom host compared to multicellular plants and animals. These results therefore demonstrate that beyond their significant morphological differences, the genomes of *Pirsonia* and oomycetes have also evolved under divergent evolutionary trajectories.

### Time-series RNA-Seq identifies infection-associated genes in *P. diadema*

To further explore the genetic contributions to *Pirsonia’s* parasitoid lifestyle, we investigated gene expression changes associated with infection progression and lifecycle transitions. To capture these changes, we sequenced bulk *Coscinodiscus radiatus* cultures at six distinct time points (t1–t6) following infection, encompassing key stages along the diatom survival curve (Fig. 3a). Principal Component Analysis (PCA) revealed a tight clustering of our biological replicates, with PC1 accounting for 32.7% of the total variance, likely reflecting temporal separation of the samples. To confirm that our RNA sequencing recapitulated infection progression, we mapped each sampled time point to a combined reference transcriptome of both the host and parasitoid, revealing an inversion in host and parasite derived reads overtime (Fig. 3b). This pattern is consistent with the *Pirsonia* lifecycle where early time points are dominated by active infections (t1-t3), while later time points are predominated by proliferating *Pirsonia* zoospores (t4-t6).

**Figure 3:**
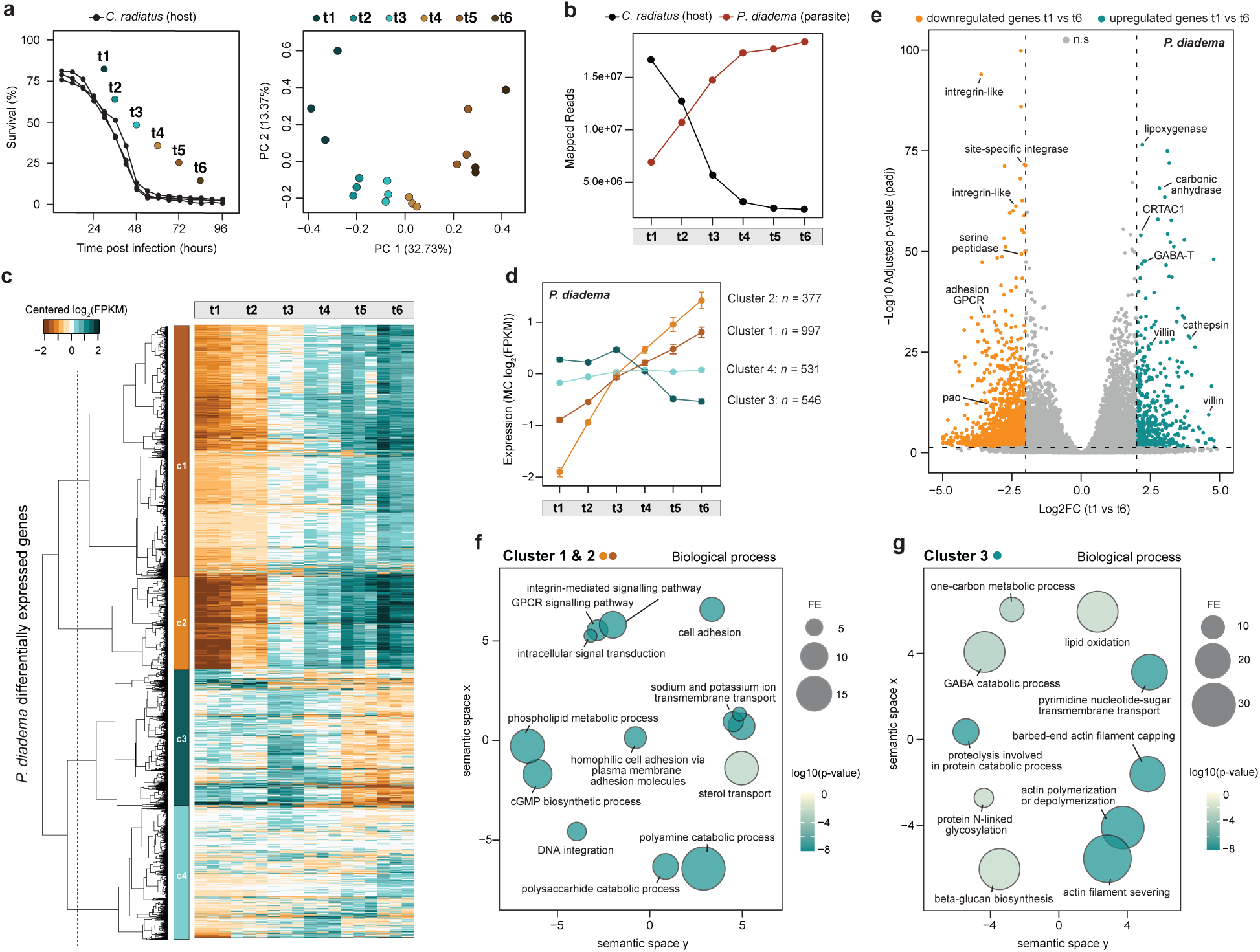
Time-resolved dual RNA-Seq of *P. diadema* infection. **a**, Survival curve and principal component analysis (PCA) of RNA-Seq samples. Line plot displaying percent host survival over time post-infection (in hours), with sampled time points for RNA-Seq labelled as t1 through t6 (Left). PCA of RNA-Seq reads, with each point representing a sample (in triplicate) and coloured according to its respective time point (Right). **b,** Line plot of mapped RNA-Seq reads for *C. radiatus* (host) and *P. diadema* (parasite) across sampled time points (t1–t6) post-infection. **c,** Clustered heatmap of 2,452 differentially expressed (DE) in *P. diadema* with a >4-fold change at *p* < 0.001 across sampled time points (t1–t6). Rows represent DE genes, and columns represent samples (and replicates). Expression values are shown as centered log₂(FPKM), where FPKM (Fragments Per Kilobase of transcript per Million mapped reads) values were log₂-transformed and mean-centered for each gene across samples. Color gradient shows relative expression changes: blues indicate upregulated genes and browns indicate downregulated genes. Hierarchical clustering (cladogram) highlights the four main expression profiles labelled as c1–c4. **d,** Line plot illustrating the average expression patterns of the four main DE gene clusters (c1–c4) in *P. diadema*. The number of genes in each cluster is displayed on the right. Expression is represented as mean-centered log₂-transformed FPKM values. Error bars indicate standard deviation. **e,** Volcano plot of significantly differentially expressed genes in t1 compared to t6. Y-axis corresponds to adjusted −log10 p-values (padj) and x-axis the log2 fold change in t1 compared to t6. Each point represents a gene, with orange points indicating genes that are downregulated in t1 relative to t6, and blue indicates genes that are upregulated in t1 relative to t6. Grey represents genes with no significant difference. **f,** Scatterplot displaying enriched GO biological process terms in *P. diadema* DE clusters 1 and 2. **g,** Scatterplot displaying enriched GO biological process terms in *P. diadema* DE clusters 3. For both plots semantic similarity was determined using REVIGO and statistical significance was assessed using two-sided permutation tests (*P* < 0.01, *n* = 10^7^).

In order to identify differentially expressed genes associated with these infection stages, we performed a differential expression analysis using DESeq2^37^. Our analysis identified 2,452 genes in *Pirsonia* with a >4-fold change (*p* < 0.001) (Fig. 3c, Table S4). Hierarchical clustering grouped these genes into four distinct co-expression clusters (Fig. 3c, Table S4), which were further categorized into three major expression patterns (Fig. 3d). Clusters 1 and 2 (1,334 genes) were highly upregulated in later timepoints (t4–t6), representing FMCs and zoospore-associated genes. In contrast, Cluster 3 (546 genes) was upregulated in early timepoints (t1–t3), suggesting an association with active infections including the formation of auxosomes and intracellular trophosomes. Cluster 4, which contained genes with mixed expression patterns across timepoints, was excluded from subsequent analyses.

To identify key genes differentiating t1 (trophosomes and auxosomes) and t6 (FMCs and zoospores), we examined the most significantly differential expressed genes and found that most were unannotated predicted proteins, consistent with the high proportion of unknown genes in the genome (Fig. 1e). Among the annotated genes, those upregulated in t1 and downregulated in t6 included cytoskeletal modifiers (e.g., villin) and protein degraders (e.g., cathepsin), while cell adhesion-related genes (e.g., integrin alpha-like genes and adhesion GPCRs) were upregulated in t6 and downregulated in t1. Overall, these findings indicate that early and late infection stages of *Pirsonia* exhibit distinct transcriptional profiles that can be linked to the two main parasitoid cell types: infective flagellates and auxosomes/trophosomes.

To further explore the functions of differentially expressed genes associated with different parasitic stages, we performed a gene ontology (GO) analysis of differential expressed genes (Fig. 3f, g). Enriched biological processes in the zoospore-associated clusters (Clusters 1 and 2) included adhesion proteins and GPCR- and integrin-mediated signalling components suggesting that *Pirsonia* flagellates may facilitate host recognition and attachment through membrane protein interactions or frustule binding. Additionally, processes related to ion homeostasis regulation, such as sodium-potassium ion transport, might contribute to host membrane remodelling, alongside the degradation of host membrane macromolecules via polysaccharide and polyamine catabolism (Fig. 3f). In contrast, genes upregulated in active infection stages (Cluster 3) were enriched in an entirely different set of biological processes such as cytoskeletal genes functioning in actin polymerization, depolymerization, severing, and capping (Fig. 3g). Other enriched functions in Cluster 3 were associated with protein degradation (e.g., proteolysis) and metabolite utilisation (e.g., one-carbon metabolism, lipid oxidation, and pyrimidine nucleotide-sugar transport). This indicates that trophosomes may use these genes to facilitate nutrient uptake from their hosts. Additionally, the enrichment of GABA catabolism suggests that *Pirsonia* may also influence host immune responses via interference of GABA receptor-mediated calcium signalling, a pathway that has been implicated in diatom stress responses^38^. Taken together, this data provides insights into the molecular mechanisms by which *Pirsonia* infects its host, revealing distinct functional strategies involving attachment and host exploitation.

### A highly expanded integrin-like gene family in *Pirsonia*

Given the enrichment of cell adhesion and integrin GO terms in both the *Pirsonia* genome annotations and upregulated genes in infectious zoospores, we examined the diversity and evolutionary history of these genes. While integrins play a fundamental role in many physiological processes in metazoans, they are not known to be broadly conserved across eukaryotes^39,40^. Analysis of the 51 differentially expressed *Pirsonia* cell adhesion genes revealed that they all contain a predominance of FG-GAP repeats and EGF domains, similar to the domains found on metazoan integrin-α and -β (Fig. 4a). These proteins vary widely in length (∼360–2,680 amino acids) and in the number of FG-GAP and EGF domains (Fig. 4b, Table S5), in contrast to canonical metazoan integrins, which are generally around 700-800 amino acids in length and contain either seven FG-GAP repeats (integrin-α) or four EGF repeats (integrin-β). Additionally, *Pirsonia* integrin-like proteins contain unique domains that are not present on metazoan integrins, such as FXa inhibition, glycoside hydrolase family 15, carbohydrate-binding C-type lectin, and sodium transport XK-related domains (Fig. 4a). Furthermore, 60% of the *Pirsonia* integrins were predicted to have signal peptides using DeepTMHMM v1^41^ and DeepLoc v2.1^42^, with 20% of proteins also containing transmembrane domains (TMD) (Fig. 4b, Table S5). Notably, some *Pirsonia* integrins possess multiple TMDs, in contrast to metazoan integrins, which function as single-pass transmembrane proteins. Collectively, these findings indicate that *Pirsonia* integrin-like genes are structurally and functionally distinct from canonical metazoan integrins.

**Figure 4:**
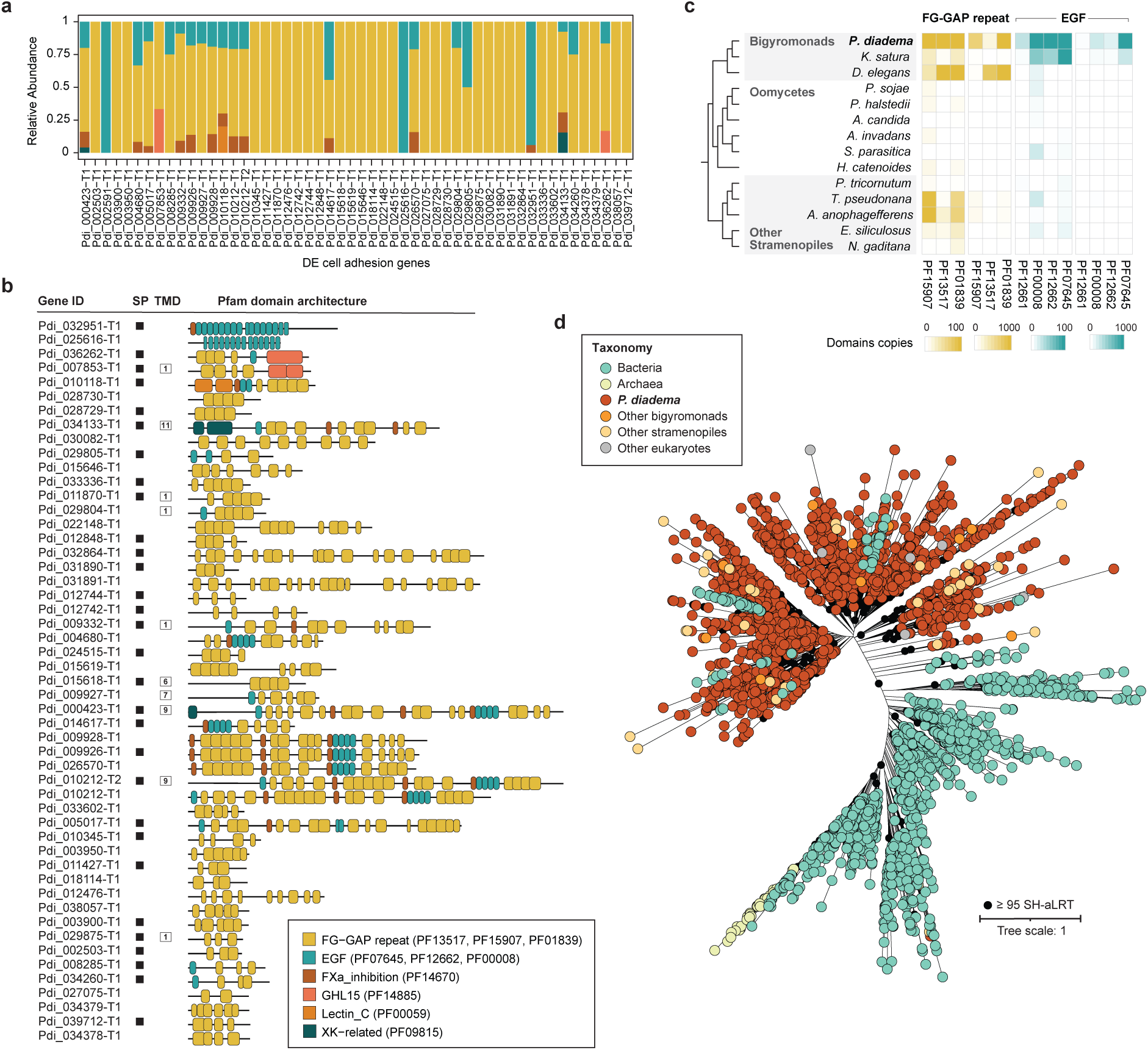
Analysis of integrin-like gene family in *P. diadema*. **a,** Relative abundance of Pfam protein domains found on DE cell adhesion genes. **b,** Protein domain architecture of DE cell adhesion genes. Presence of a signal peptide is denoted by a black square. Transmembrane domains (TMD) are shown in white squares with the number of TMDs written inside. For both a, and b, domains are coloured by Pfam domain (with FG-GAP and EGF terms collapsed). Colour key is shown below. **c,** Heatmap of number of FG-GAP (yellow) and EGF (blue) domains found in a reference set of stramenopile proteomes. Number of domains per proteome is shown in two scales: up to 100 and 1000. Pfam domain IDs are shown below. **d,** Maximum likelihood phylogeny of FG_GAP_3 domain constructed in IQ-Tree. Support was obtained using 1,000 SH-like approximate likelihood ratio tests (SH-aLRT). Bootstraps greater than 95% are shown as black circles. Tips are collapsed and coloured by species. Full tree can be viewed at https://itol.embl.de/shared/1K4uJeMydJIHt.

To investigate their evolutionary history, we performed HMM searches for FG-GAP repeat- and EGF domain-containing proteins across diverse stramenopiles (Fig. 4c). Although we identified proteins containing these domains in oomycetes and other stramenopiles, they are significantly more abundant in *Pirsonia*, which has 445 integrin-like genes annotated in the genome, containing thousands of FG-GAP and EGF domains (Fig. 4c). The presence of introns within these genes indicates they are unlikely to originate from bacterial contamination (Fig. 4c, Table S5). Notably, bigyromonads like *K. satura*, the sister group to *Pirsonia,* also contains many EGF domain-containing proteins but fewer FG-GAP domain proteins, whereas *D. elegans,* another related bigyromonads species, shows the opposite pattern. This suggests that genes encoding these domains were present in the last common ancestor of stramenopiles and underwent significant expansion in bigyromonads, particularly in *Pirsonia*, potentially as an adaptation to diatom parasitism.

To better understand the evolutionary history of FG-GAP-like repeat-containing proteins, we expanded our search to a balanced set of taxonomically diverse reference proteomes across the tree of life (see Materials and Methods) and constructed a maximum likelihood phylogeny (Fig. 4d). We identified orthologs primarily from diverse bacterial lineages as well as photosynthetic ochrophytes (stramenopiles), and various other eukaryotic groups; cryptomonads (*Guillardia*), haptophytes (*Diacronema*), and green algae (*Ostreobium*). The eukaryotic sequences formed a distinct clade, separate but related to a large assemblage of bacterial sequences. Given the currently available genomic datasets, this topology may reflect either an ancient eukaryotic origin followed by extensive gene loss across most eukaryotic lineages, or a horizontal gene transfer from bacteria followed by a lineage-specific expansion in *Pirsonia*. Resolving these scenarios will require a deeper genomic sampling of undersampled eukaryotic lineages, as well as functional characterization studies of these FG-GAP-like proteins.

### Cytoskeletal drugs impair *P. diadema* infection

In metazoans, integrins serve as linkers between the extracellular matrix and the intracellular actin cytoskeleton^40^. Given the convergence in protein domains and transmembrane localization of *Pirsonia* FG-GAP- and EGF-domain containing genes with canonical integrins, along with the enrichment of actin-related functions in auxosomes/trophosomes (Fig. 3g), we hypothesized that *Pirsonia*’s infection strategy may be dependent on cytoskeletal dynamics.

To test this, we assessed the importance of actin during infection by treating infected diatoms with three actin-disrupting agents and monitored infection progression via using our live-cell time-series high-content imaging approach (Fig. 5a, Fig. S3). Cytochalasin D, which caps actin filaments and blocks polymerization, latrunculin B, which sequesters actin monomers, and jasplakinolide, which stabilizes filaments, all impaired infection under standard working concentrations. Treated cells showed complete diatom survival (Fig. 5a), no successful infections (Fig. 5b), and stable zoospore numbers (Fig. 5c), indicating that dynamic actin remodelling is essential for *Pirsonia* infection progression but not because it causes zoospore death. For comparison, we also treated cells with taxol (paclitaxel), a microtubule-stabilizing drug, that we expected to interfere with zoospore function as they rely on microtubule-based flagella motility^43,44^. Consistent with this, taxol treatment also impaired infection (Fig. S3), suggesting that both cytoskeletal systems are required for infection progression. While microtubules appear essential for zoospore function, we hypothesized that actin may be specifically required for host attachment and trophosomes formation. To investigate this, we more closely examined parasite attachment using SEM (Fig. 5d) and identified clear pseudopod-like extensions at the contact sites between *Pirsonia* auxosomes and diatoms. Although genetic manipulation will ultimately be required to test these hypotheses, our findings support a model in which actin facilitates auxosome adhesion and intracellular trophosome release. This suggests that dynamic remodelling of the actin cytoskeleton may drive *Pirsonia* infection progression via actin-filled pseudopods.

**Figure 5:**
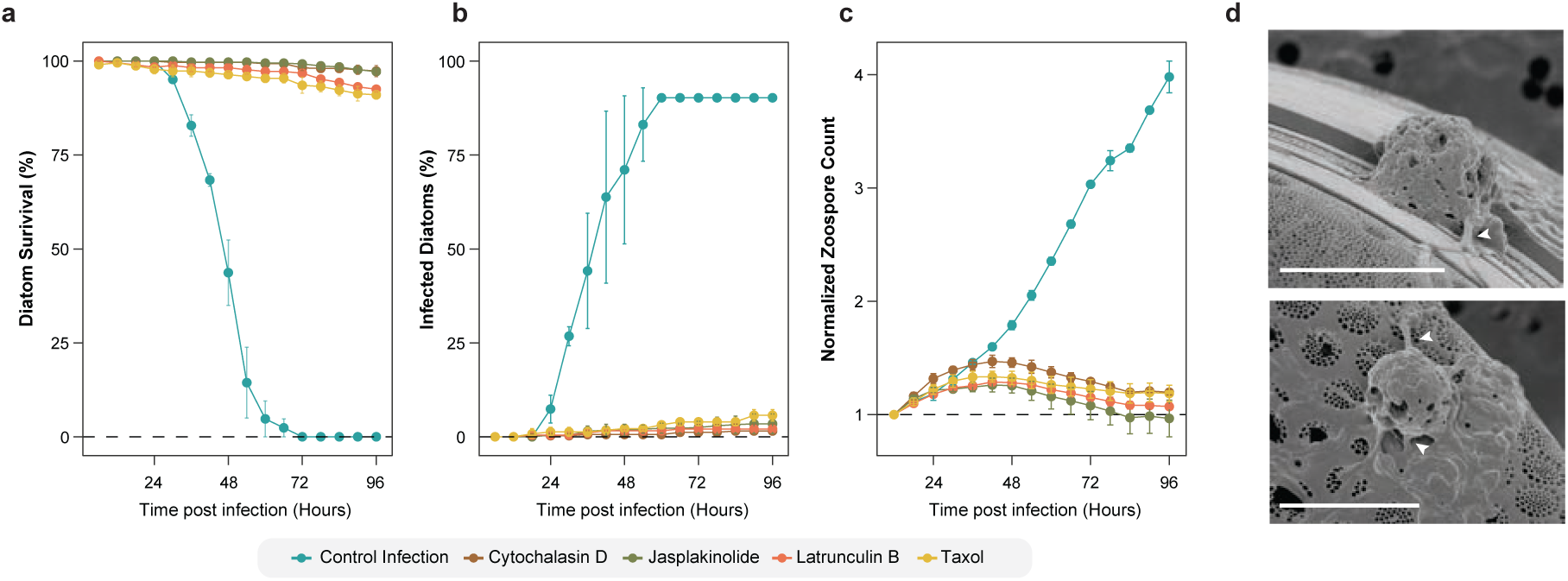
High-content screening (HCS) of *P. diadema* infecting *C. granii* under various cytoskeletal drug treatments. Each condition was performed in triplicate; error bars represent standard deviation. **a,** Percentage of diatom survival over time post-infection, with survival defined as CTCF remaining above 50% of the averaged baseline. **b,** Percentage of diatoms infected over time, calculated by a 15% increase in cell area. **c,** Zoospore counts normalized to initial values per well, measured throughout the infection. **d,** Scanning electron microscopy (SEM) of *Pirsonia*’s attachment site on diatom frustule, white arrows indicate pseudopodia. Scale bars = 5 μm.

## DISCUSSION

Here, we establish a model pathosystem for investigating phytoplankton–parasite interactions, composed of the globally distributed, bloom-forming, environmentally important diatom, *Coscinodiscus* and its parasitoid *Pirsonia*. In combination with the development of phenotyping assays based on high-throughput live-cell imaging, this system offers a unique platform to dissect the molecular and cellular mechanisms underlying a marine host–parasite interaction. The rapidity of the *Pirsonia* life cycle and its capacity to cause total diatom population collapse in culture highlights the ecological importance of this interaction and the capacity for microbial parasites to shape biogeochemical nutrient cycling in a similar manner to marine viruses^45^.

The development of the first genome assembly of *Pirsonia diadema* (Fig. 2), a representative of the largely free-living bigyromonads, and the identification of infection-associated genes using RNA-Seq, provide a unique comparative system for examining parasitic evolution. The identification of integrin-like proteins, as a candidate gene family of variable surface proteins in *Pirsonia* reveals the presence of a unique yet convergent hallmark of parasite genomes^46^. These uncharacterized FG-GAP proteins may function in surface adhesion or recognition of host-derived ligands, reminiscent of strategies employed by other eukaryotic parasites such as *Plasmodium*^47^, trypanosomes^48^, and oomycetes^49^, although other parasitic species have co-opted alternative gene families for this function. Integrin expansion in *Pirsonia* suggests a lineage-specific adaptation to diatom parasitism, possibly mediating attachment to the diatom frustule or facilitating entry into the host cytoplasm. Together, these findings establish a genomic framework for dissecting virulence mechanisms in a key marine parasite and emphasize the importance of surface protein diversification in parasitic evolution across the tree of eukaryotes.

The susceptibility of *Pirsonia* infection to actin inhibitors suggests that auxosomes and trophosome formation may be mediated through an actin-dependent mechanism, as hypothesized in early studies^50^. This strategy appears to have evolved in parallel across diverse parasitic lineages. For example, apicomplexans use the actin-myosin-based glideosome for host invasion^51^, while the chytrid fungus *Batrachochytrium*, which infects amphibian skin, dynamically remodels its actin cytoskeleton during infection^52^. A pseudopodia-based adhesion mechanism could also play an ecological role in facilitating a firm attachment to the host in the turbulent surface waters inhabited by *Pirsonia*^24^. Notably, a recent study taxonomically characterizing relatives of *Pirsonia* show their close relatives to be numerous predatory species that engulf and digest entire eukaryotic prey using pseudopodia^31^. However, the formation of auxosomes and trophosomes remain unique to *Pirsonia* parasitoids. This suggests that while phagocytosis and pseudopodia are likely ancestral traits linked to predation, *Pirsonia* has co-opted and repurposed these mechanisms to serve a parasitic lifestyle. Intriguingly, this trait blurs the distinction between parasitism and predation, positioning *Pirsonia* as a key group for studying evolutionary shifts in trophic strategies and the origins of parasitism. Expanded genomic sampling of more *Pirsonia* species and their free-living relatives along with detailed cell biology will be necessary to uncover the molecular innovations underpinning this shift.

Another key question about this interaction is whether the diatom mounts an immune response and whether, under natural conditions, parasitoids and their hosts coexist, similar to the coevolutionary arms race observed between marine viruses and their hosts^53^. However, our bulk RNA-Seq approach was unable to resolve the host’s transcriptional response to *Pirsonia* infection beyond loss, as we primarily captured the diatom’s death response (Fig. S2, Table S6). This may reflect differences in immune pressure: parasites infecting multicellular hosts, such as the plant-infecting oomycete *Phytophthora sojae*, face intense immunological challenges and have evolved specialized effectors, such as decoy effectors produced by gene duplication which neutralize host defenses^54^. In contrast, the apparent lack of small effectors in *Pirsonia* suggests that its unicellular diatom host exerts a comparatively reduced immune pressure, allowing retention of a broader, less specialized secretome consistent with the large, gene-rich genome we observe. However, the absence of detectable host immune responses in our bulk RNA-Seq could also stem from methodological limitations, as subtle or transient responses may be masked in population-level analyses. To address this, single-cell RNA sequencing approaches, recently demonstrated to reveal nuanced host-virus interactions in planktonic systems^55^, provide a promising avenue to uncover potential early host-specific responses and Red Queen dynamics driving host–parasitoid coevolution in natural marine environments.

Finally, these interactions occur within complex microbial communities, and growing evidence highlights that diatoms engage in diverse biotic relationships with bacteria and other microorganisms^56,57^. Therefore, incorporating more microbial players into this model will be essential to fully unravel the multifaceted nature of host-parasite dynamics in marine ecosystems. While the ecological importance of parasites in oceanic food webs is increasingly recognized^15,58,59^, key aspects of parasite-mediated phytoplankton mortality and its consequences remain largely unexplored. By establishing a new model pathosystem, our study provides a critical foundation for elucidating the molecular and ecological mechanisms underlying marine parasitism.

## MATERIALS AND METHODS

### Sampling and cultivation of host and parasite

*Coscinodiscus radiatus* and *Coscinodiscus granii* were isolated from marine plankton tows carried out at Station L4 (50°20′0.093″N, 4°08′0.915″W) located southwest of Plymouth in the Western English Channel. To establish cultures, *Coscinodiscus* single cells were manually isolated using a microcapillary pipette, washed multiple times in autoclaved artificial seawater, and propagated in f/2 + Si media (Guillard’s medium for diatoms). *Pirsonia diadema* was isolated from the net-tow sample at the Yongho Bay (35°08′00″N, 129°06′55″E) of Busan, Republic of Korea*. P. diadema* was cultivated by inoculating 20 mL of *C. radiatus* with 1 mL from a 4-day old, infected culture. Both host and parasite cultures were maintained in a 12-hour light-dark cycle at 21°C.

### Scanning electron and confocal microscopy

For SEM, *C. radiatus* infected with *P. diadema* were fixed by adding 25% Glutaraldehyde in artificial seawater (ASW) directly into a 3-day old 20 mL culture followed by a 4-hour incubation at 4°C. Fixed cells were gently filtered onto a Whatman® Nucleopore™ Track-Etched Membrane with a pore size of 0.8 µm. Cells on the filter were gently washed three times with Milli-Q water and dehydrated in a graded ethanol series (30, 50, 70, 80, 90 and 99%) with a 15-minute incubation at each step. Cells were chemically dried with a 10-minute incubation followed by an overnight incubation in 100% hexamethyldisilane (HMDS). The filter was critical point dried with CO_2_ and sputter coated prior to imaging.

Confocal microscopy was performed by concentrating 40 mL of infected diatoms via gravity filtration using a 40 μm pluriStrainer (pluriSelect). Cells were resuspended in 0.5 mL of ASW, and a 100 μL drop was placed on a poly-L-lysine–coated Poly-Prep glass slide (Sigma-Aldrich) for 1 hour to allow sedimentation and attachment. Excess ASW was removed by gently tipping the slide and blotting the edges with filter paper. Cells were fixed in 250 μL of 4% (w/v) paraformaldehyde in ASW for 15 minutes at room temperature. The fixative was replaced with 250 μL ASW for a 1-minute wash. Cells were then permeabilized in 250 μL PBS with 0.1% (v/v) Triton X-100 in ASW for 5 minutes at room temperature. After a second wash, Hoechst 33342 (Sigma-Aldrich, Cat. No. H3570) was added at a 1:500 (v/v) dilution for 60 minutes at room temperature. Following two additional washes, 7 μL of ASW was added to the slide, which was then covered with a coverslip and sealed with transparent nail polish to prevent evaporation. Imaging was performed on a Zeiss LSM-780 confocal microscope using x20 and Ph3 x60 oil objectives. Exposure settings were kept constant, and images were acquired with ZEISS ZEN software and processed using Fiji (ImageJ v1.54)^28^.

### Environmental survey analysis

rDNA 18S V9 metabarcoding tables (Swarms) for Tara Oceans Expedition (2009-2013), including Tara Polar Circle Expedition (2013), were obtained from Zenodo (https://zenodo.org/records/7236051)^28^. Swarm taxonomy tables were queried for *Pirsonia* and *Coscinodiscus* operational taxonomic units (OTUs). Samples containing fewer than three OTUs of either *Pirsonia* or *Coscinodiscus* were excluded. Geographical distributions (Fig. 1d) and OTU abundances (Fig. S1) were visualized using custom python scripts.

### Timeseries high-content screening (HCS), drug assays, and image analysis

For life cycle quantification, *C. radiatus* was added to a 96-well plate and inoculated with *P. diadema* zoospores. Images were captured every 6 hours for 96 hours (16 time points) using an ImageXpress Pico Automated Cell Imaging System (Molecular Devices) at 10× magnification in both brightfield and Cy5 channels (absorbance 630/40 nm; emission 695/45 nm). Images were stitched into a composite “tiled” image of the entire well using the CellReporterXpress software.

For cytoskeletal drug assays, *C. granii* was added to a 96-well plate and inoculated with *P. diadema* zoospores and imaged under different drug conditions using a Zeiss Cell Discoverer 7 microscope. Brightfield and Cy5 channels were used to capture images every 6 hours for 96 hours (16 time points) at 21 °C with a 5× objective. Images were stitched using ZEN Imaging Software. All drugs were diluted in DMSO to a final concentration of 0.3% in all wells. The drug conditions tested were cytochalasin D (5 µM), latrunculin B (5 µM), jasplakinolide (3 µM), and taxol (5 µM). All treatments were performed in triplicate (i.e., three wells per condition) alongside DMSO only controls.

Image analysis was conducted in Fiji (ImageJ v1.54)^28^ using the Labkit plugin for supervised segmentation. Measurements including integrated density (IntDen), mean fluorescence, and cell area, were extracted per diatom cell across all time points. The same was done for zoospores. Corrected Total Cell Fluorescence (CTCF) was calculated as: CTCF = IntDen − (area × background). Outliers in CTCF and area were removed using an interquartile range filter. Cells were tracked based on their positional coordinates and only cells tracked across all time points and alive at the start of the experiment were included in downstream analysis. Survival curves defined live cells as those maintaining CTCF above 50% of the well-averaged baseline. Infection dynamics were quantified by tracking individual cells and classifying them as infected if their area increased by ≥15% relative to baseline. To reduce false positives from transient shape changes, size trajectories were cross-validated against the final size. Zoospore counts were filtered by area to exclude debris, averaged across replicates, and normalized to the first time point. All analyses and visualizations were performed in R v2023.12.1+402 (ggplot2, dplyr) and Python v 3.13.3.

### PacBio HiFi genome sequencing and assembly

One litre of dense *C. radiatus* cultures were infected with *P. diadema* zoospores. At four days post-infection, the cultures were filtered through a cell strainer (pore size = 40µm) to exclude the diatoms (empty frustules). The resulting sample, composed of *P. diadema* flagellates, was then filtered onto a Whatman® Nuclepore™ Track-Etched Membrane (pore size = 0.8 µm) to exclude bacterial cells. *P. diadema* cells were scraped off the filter and used for the high-molecular weight DNA extraction using the Qiagen MagAttract HMW DNA Kit (Cat. No. 67563). DNA Fragmentation was performed using g-TUBEs (Covaris) with a target size of 15 kbp. Library preparation was performed using the SMRTbell Express TPK 2.0 and SMRTbell Enzyme Clean-up kit v1 (Pacific Biosciences). Sequencing was performed on a Sequel IIe PacBio™ machine. The “Run Design Application” was set to HiFi Reads with default settings. The resulting sequencing library comprised 739,524 reads with lengths up to 55,211 bases. Reads were filtered using “HiFiAdapterFilt” to remove adapter-contaminated reads, resulting in 605,547 reads^60^. Bacterial contamination was removed using Kraken2^61^ with the “Standard plus” RefSeq database that includes protozoa & fungi. The cleaned reads were assembled using HiFiasm v0.19.7^29^. BLASTn^62^ searches were carried against NCBI nt database^63^ to remove any bacterial and diatom contamination using BlobTools2^64^. The cleaned reads were re-assembled resulting in an assembly of 5,155 contigs and assembly size of 171Mb (N50 = 42.1Kb). BUSCO v6.0.0 (Benchmarking Universal Single Copy Orthologues)^65^ was used to assess the quality and completeness of the genome assembly relative to the stramenopile_odb10 database. Raw reads were deposited in NCBI SRA SRR34520686.

### *Pirsonia diadema* genome annotation

Annotation of the *P. diadema* genome assembly was performed using Funannotate v1.8.15 (https://github.com/nextgenusfs/funannotate). *P. diadema* transcripts (see below for details) were mapped to the genome using minimap2^66^ to generate homology-based gene predictions in “PASA” (Program to Assemble Spliced Alignment)^67^. *Ab initio* gene predictions were also performed using a combination of AUGUSTUS^68^, GeneMark-ES^69^, SNAP (https://github.com/KorfLab/SNAP), BUSCO^65^ and GlimmerHMM^70^. tRNAs were predicted using tRNAscan-SE^71^. Evidence Modeler was used to consolidate the gene predictions and output consensus gene models^67^. The final gene set were functionally annotated using several databases (EGGNOG^72^, MEROPS^73^, InterProScan5^74^, CAZy^75^, PFAM^76^ and UniProt^77^). Additionally, some of the putatively mis-predicted genes were manually removed based on the following criteria: i) over-represented ortholog copies ii) lack of support from RNA-Seq data, ii) lack of homology to other proteins, and iii) lack of a predicted domain inside the protein. If all three of these conditions were met, the genes were excluded. The mitochondrial genome was assembled using MitoHifi^78^ and annotated using MFannot (https://megasun.bch.umontreal.ca/apps/mfannot/) using the standard genetic code. Gene annotations were deposited Mendeley Data DOI: 10.17632/xc89mvhxpg.

### Comparative genomics and phylogenomic tree construction

Pseudofungi proteomes were compiled from UniProt^77^, in addition to transcriptomes retrieved from Cho et al., 2022^32^. Orthogroup inference was performed using OrthoFinder v2.5.5^35^ and upset plots were generated using the ComplexUpset package^79^ in RStudio v2023.12.1+402. GO enrichments were assessed by comparing the frequency of individual GO terms in *P. diadema*-specific genes to the whole proteome (i.e. the background). Significance was assessed using permutation tests with test distributions produced by generating 100,000 randomized samples, equal in size to the test-set, by randomly selecting transcripts from the transcriptome without replacement. The highly enriched GO-terms (*p* < 0.001) were summarized and visualized using REVIGO^80^.

For phylogenomic analyses, *P. diadema* proteins were searched against a set of 263 genes, previously used for eukaryote-wide phylogenetic analyses^23,32^ using BLASTp^62^. Resulting hits were filtered using an e-value threshold of 1e-20 and a minimum query coverage of 50%. Individual alignments were produced using mafft^81^ and trimmed using trimAl^82^ (gap threshold = 0.6). Single gene trees were then constructed using IQ-TREE^83^ and manually inspected to remove paralogs and contaminants. The final cleaned gene-sets were filtered so that they contained a maximum of 60% missing positions and concatenated. The resulting concatenated alignment consisted of 241 genes spanning 79, 399 amino acid positions from 45 taxa. A maximum likelihood phylogeny was generated from the concatenated alignment in IQ-TREE^83^ using the heterogenous mixture LG+C40+F+G4 model. Statistical support was assessed using 100 non-parametric bootstraps and 1,000 SH-aLRT support using the Q.yeast+F+I+R6 model as inferred by ModelFinder^84^. Phylogenomics statistics can be found in Table S1 and alignments and phylogenies were deposited Mendeley Data DOI: 10.17632/xc89mvhxpg.

### Host and parasite reference transcriptomics

*C. radiatus* cells were collected during both the light cycle (AM) and night cycle (PM). 150mL of dense *C. radiatus* cultures were filtered and washed on a cell strainer (pore size = 40 µm) and filtered on a membrane filter (pore size =10 µm). The filter was directly immersed in 1.5 mL of Invitrogen TRIzol reagent and incubated at room temperature for 10 minutes followed by a 60°C incubation for 3 minutes. The samples were centrifuged at 13,000 g for one minute to pellet the empty diatom frustules. *P. diadema* cells were harvested on day four from a 400 mL infected *C. radiatus* culture. The culture was filtered on a cell strainer (pore size = 40 µm) to exclude diatoms. The flow-through was then filtered (pore size = 10 µm) to exclude smaller diatom cells and broken frustules. The filtrate was centrifuged at 4,000 g for 20 min and the pellet was resuspended in 300 µL of Invitrogen TRIzol reagent. For both *C. radiatus* and *P. diadema*, RNA extraction and purification was performed using the Direct-zol RNA Miniprep kit (ZymoResearch, Irvine, CA). RNA concentration and quality was assessed with the Bioanalyzer RNA 6000 Nano assay kit and Qubit RNA Broad Range assay kit.

Illumina sequencing libraries were prepared using the Novogene NGS RNA Library Prep Set (PT042) and sequenced on the Illumina Novaseq 6000 platform. The resulting raw reads were trimmed using Cutadapt^85^ to remove adapter sequences. Kraken 2^61^ was used to taxonomically classify the transcripts and remove bacterial contamination. The cleaned reads were assembled using rnaSPAdes^86^. Protein prediction were performed using TransDecoder (https://github.com/TransDecoder/TransDecoder).The proteomes underwent an additional round of bacterial contamination removal (and diatom contamination removal in the case of *P. diadema*) using Blob Toolkit^64^ and DIAMOND^87^ searches against the Swiss-Prot database^77^. The AM and PM *C. radiatus* datasets were merged for downstream analyses. Raw reads were deposited in the NCBI BioProject PRJNA1291136 and PRJNA1290282 and predicted proteins in Mendeley Data DOI: 10.17632/xc89mvhxpg.

### Time-resolved dual bulk RNA-sequencing

*C. radiatus* cultures were infected with 40 mL of *P. diadema* zoospores (in three replicates). 24-hours post infection, 50 mL was sub-sampled from the infected culture and filtered on a membrane filter (pore size = 0.8 µm). One mL of TRIzol reagent (Invitrogen) was added directly to the cells onto the filter and incubated at 60°C for two minutes. Samples were centrifuged at 13,000g for one minute to pellet diatom frustules and the downstream RNA extraction was carried out using the Direct-zol RNA Miniprep kit (ZymoResearch, Irvine, CA). The same process was repeated every 12 hours until the end of the infection cycle at 96 hours (for a total of 6 time points). The quality of the RNA was assessed with the Bioanalyzer RNA 6000 Nano assay kit and Qubit RNA Broad Range assay kit. Illumina sequencing libraries were prepared using poly-A selection Novogene NGS RNA Library Prep Set (PT042) and sequenced on the Illumina Novaseq 6000 platform. Raw reads were deposited in NCBI BioProject PRJNA1292109.

### Differential Expression (DE) and Gene Ontology (GO) enrichment analysis

The raw Illumina paired-end sequencing reads were trimmed for adaptors and low-quality sequences using Cutadapt^85^. The expression of transcripts across samples (t1-t6) and replicates was quantified using Salmon^88^ by quasi-mapping the reads to combined reference proteome of both host and parasite. Trinity^89^ was used to perform cross-sample normalization (measured as transcripts per million (TPM)) and to generate a transcript count matrix. DESeq2^37^, implemented in RStudio v2023.12.1+402, was used to detect DE transcripts and extract DE genes that were at least four-fold differentially expressed at a significance of ≤ 0.001 in any of the pairwise sample comparisons. DE genes were hierarchical clustered with hclust in R using the ward.D2 method. For GO enrichment analysis of the DE genes, we first annotated our reference *C. radiatus* and *P. diadema* transcriptomes with GO terms using Wei2GO. This involved DIAMOND^87^ and HMMScan (http://hmmer.org/) searches against the UniProtKB ^77^ and Pfam^76^ databases and using a weighing algorithm to calculate a score for the Gene Ontology annotations. GO enrichment was assessed by comparing the frequency of individual GO terms in the DE genes to the whole proteome (i.e. the background). Significance was assessed using permutation tests with test distributions produced by generating 100,000 randomized samples, equal in size to the test-set, by randomly selecting transcripts from the transcriptome without replacement. The highly enriched GO-terms (*p* < 0.001) were summarized and visualized using REVIGO^80^.

### Domain analysis and phylogeny of integrin-like proteins

To identify the protein domains present on *P. diadema* cell adhesion genes, profile Hidden Markov Model (HMM) searches were performed using the Pfam^76^ database and HMMER^91^ (e-value threshold = 1e-5 and domain e-value threshold = 1e-5). The resulting domains were used to conduct HMM searches against our curated set of reference Pseudofungi proteomes (see above). The resulting hits were extracted and used to generate a domain heatmap (Figure 4b). Signal peptides and transmembrane domains were predicted using DeepTMHMM v1^41^and DeepLoc v2.1^42^.

Next, we compiled a diverse reference dataset comprising proteomes from UniProt^77^. To taxonomically balance the dataset, we selected the best two proteomes per genus for eukaryotes (n = 141) based on BUSCO^65^ completeness. For metazoans, fungi, and embryophytes, more strict taxonomic criteria were set by selecting the best proteome per phyla for metazoans (n = 17), the best proteome per class for fungi (max two per phylum, n = 12), and the best proteome per order for embryophytes (max three per class, n = 11). For bacteria, the best proteome per species up to 2 per genus (n = 4,833) was selected, and for archaea, the best proteome per species up to 2 per genus (n = 310). For viruses, the best proteome per species up to 1 species was selected based on protein count (n = 11,104). Each individual proteome was then clustered at 99% sequence identity using CD-HIT^92^.

The longest protein domain (PF13517: FG-GAP_3) on the integrin-like genes was used as a query for a BLASTp^62^ search against this curated reference dataset. The resulting hits were filtered at an e-value threshold of 1e-25 and clustered at 97% sequence identity using CD-HIT^92^. The hits were aligned using MAFFT^81^ and trimmed with a gap threshold of 60% using trimAl^82^. The alignment was filtered to remove sequences that contained less than 50% of amino acid sites. ModelFinder^84^ was used to identify the best-fit evolutionary model according to the Bayesian Information Criterion, and a maximum likelihood phylogeny was constructed in IQ-TREE^83^ with support from 1,000 SH-like approximate likelihood ratio tests (SH-aLRT) and 1,000 ultrafast bootstraps. Eukaryotic operational taxonomic units (OTUs) forming clades with single species branching within prokaryotes and vice versa for prokaryotic sequences were excluded. Tree editing and visualization were done in iTOL v6^93^. Alignments and phylogenies were deposited in Mendeley Data DOI: 10.17632/xc89mvhxpg.

## ACKNOWLEDGEMENTS

We thank Hyewon Kim for *Pirsonia* culture establishment; Theodosios Kyriakou at the Oxford Genomics Centre (OGC) for assistance with PacBio sequencing; Jennifer Holter for support with SEM; Guy Leonard for bioinformatics assistance; Alberto Moreno Cencerrado at VBC BioOptics for help with high-content screening (HCS); Estelle Kilias for laboratory support and culture maintenance; and Magnus Nordborg for providing resources and mentorship.

## AUTHOR CONTRIBUTIONS

Conceptualization: V.M, T.A.R. Methodology: V.M., N.A.T.I., T.A.R. Investigation: V.M., N.A.T.I., L.J.G., E.A., M.C., S.K. Supervision: T.A.R, U.T. Writing—original draft: V.M., T.A.R. Writing—review & editing: all authors.

## SUPPLEMENTAL INFORMATION

### Supplemental Figures

Figure S1: Tara Oceans SWARM abundance (maps)

Figure S2: Diatom RNA-Seq heatmap

Figure S3: HCS images from drug treatments

### Supplemental Tables

Table S1: Tara Oceans Swarms and sample info

Table S2: Phylogenomics tree statistics

Table S3: OrthoFinder orthogroups

Table S4: DE genes: *Pirsonia diadema*

Table S5: Pirsonia integrin genes: length, pfam domains, localization/TMD, introns

Table S6: DE genes: *Coscinodiscus radiatus*

### Supplemental Video

Supplemental video 1: HCS of *Pirsonia* infection cycle

