## Supplementary figures and images for "Phagocytosis repurposed: infection strategies in a global marine diatom-parasite interaction"

### Fig. S1

#### Figure S1

Tara Oceans Expedition (2009-2013), including Tara Polar Circle Expedition (2013)

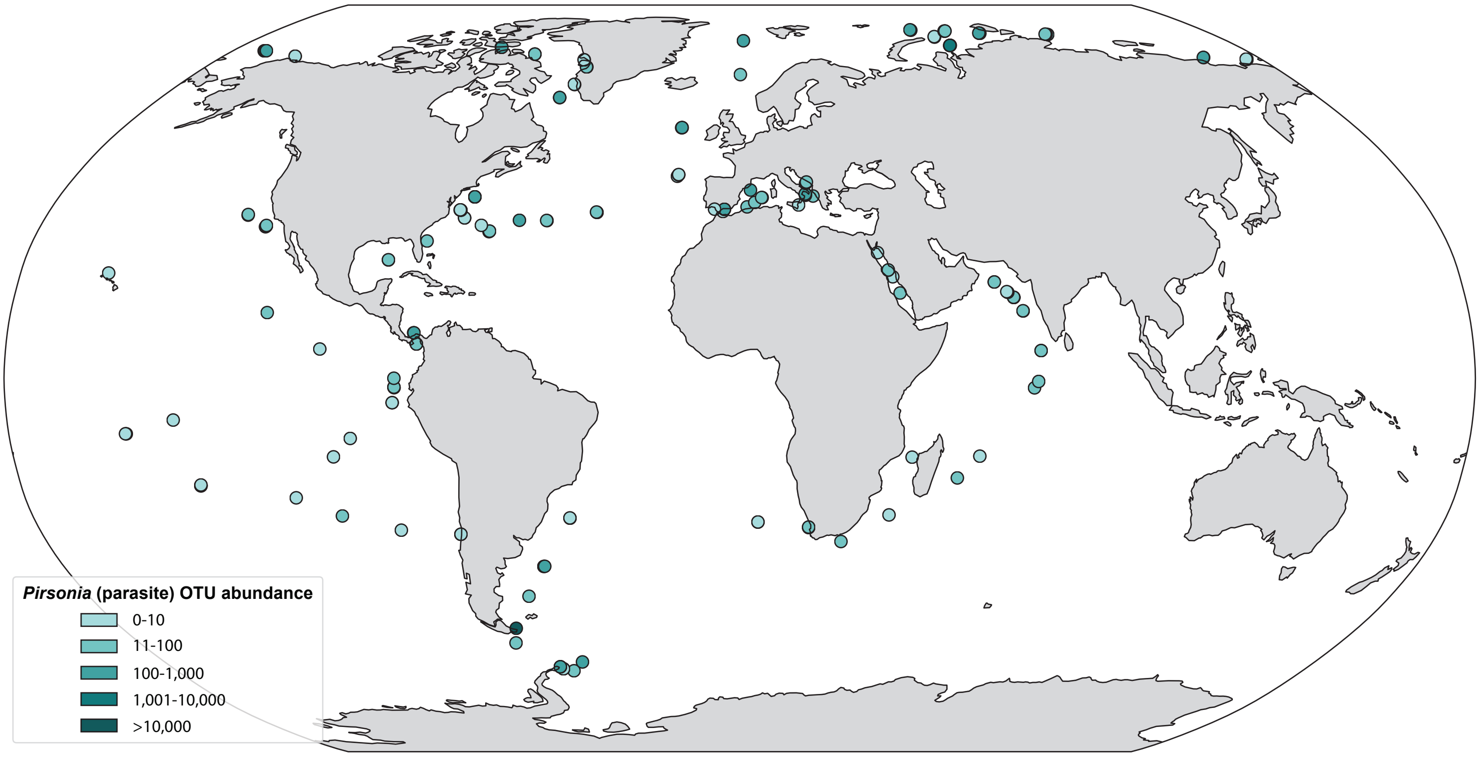


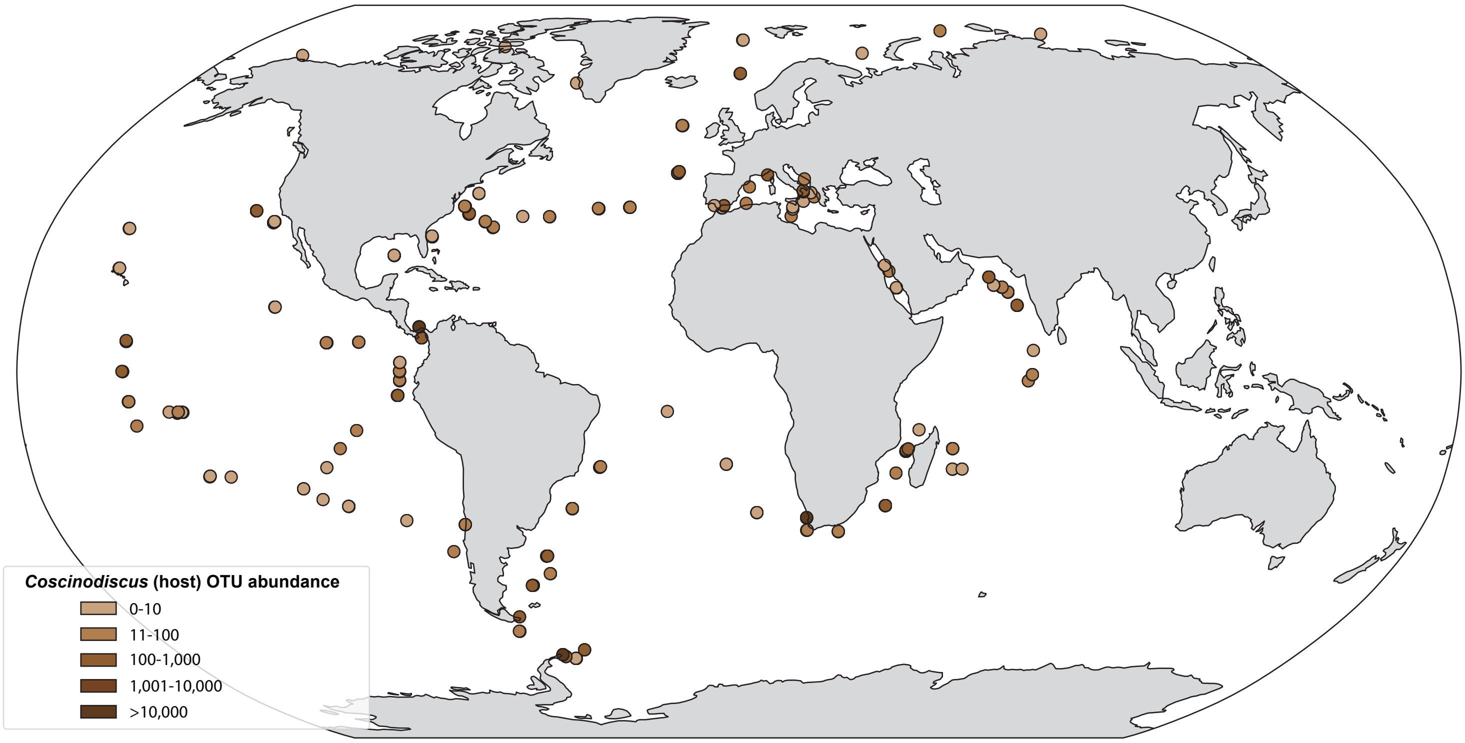

### Fig. S2

##### Figure S2 : RNA-Seq DE analysis for *Coscinodiscus radiatus* (host)

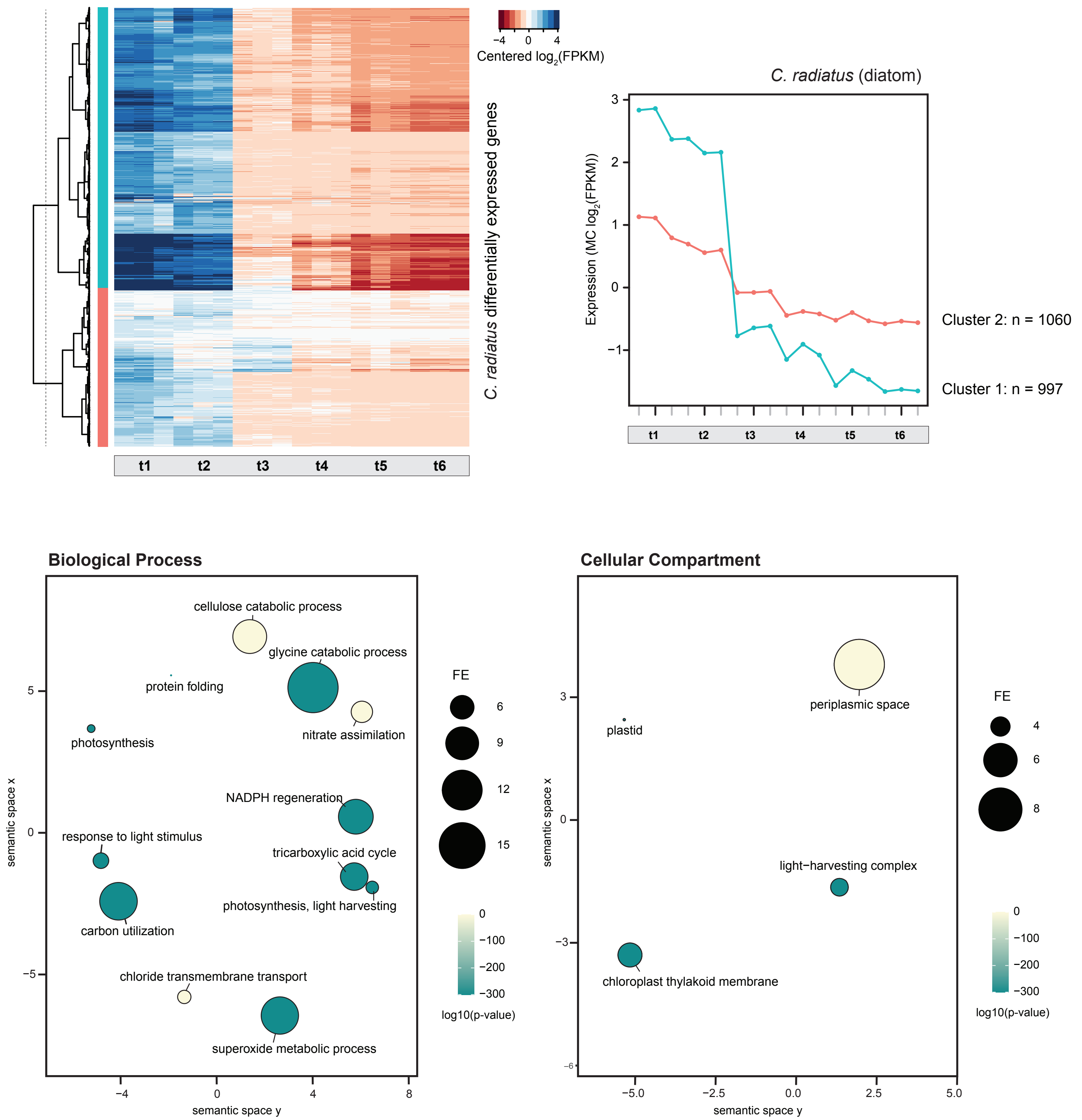

### Fig. S3

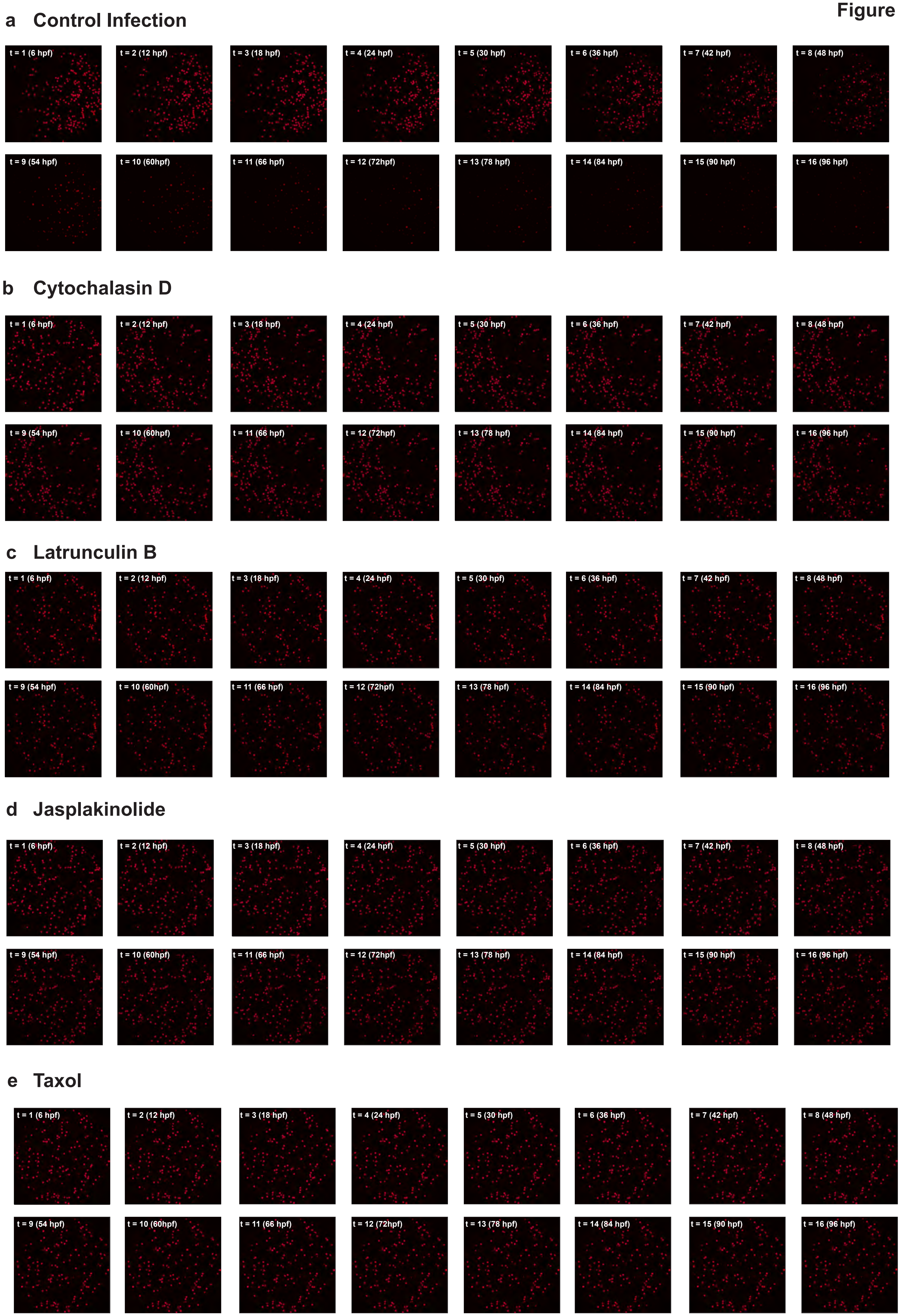
